# Functional Profiling of Kiwifruit Phyllosphere Bacteria: Copper Resistance and Biocontrol Potential as a Foundation for Microbiome-Informed Strategies

**DOI:** 10.64898/2026.01.11.698881

**Authors:** Vinicius Casais, Joana Pereira, Eva Garcia, Catarina Coelho, Daniela Figueira, Aitana Ares, Igor Tiago, Joana Costa

## Abstract

Bacterial canker, caused by *Pseudomonas syringae* pv. *actinidiae* (Psa) is a major threat to global kiwifruit production. Copper-based bactericides remain widely used, but increasing resistance underscores the need for alternative strategies. Understanding the functional capabilities of phyllosphere bacteria under copper pressure is critical for designing sustainable alternatives. This study provides a culture-based functional inventory of bacteria associated with *Actinidia chinensis* var. *deliciosa* leaves from Portuguese orchards under long-term copper management, aiming to identify native taxa with traits relevant to plant health and resilience. A total of 1,058 isolates were recovered and grouped into 261 RAPD clusters, representing 58 species across 29 genera. Representative strains were screened for plant growth-promoting (PGP) traits (IAA, siderophore production, phosphate solubilization, ammonia production), copper tolerance, and in vitro antagonism against Psa. Copper resistance was widespread (53.3% of isolates with MIC ≥0.8 mM), including the first evidence of a highly copper-resistant PSA in Portuguese kiwifruit orchards and an exceptionally resistant non-pathogenic strain (*Erwinia iniecta*, MIC 2.8 mM). A subset of 25 isolates combined all PGP traits. In parallel, several isolates exhibited antagonism against Psa, including *Bacillus pumilus*, which consistently inhibited pathogen growth. Notably, some antagonistic isolates overlapped with the multifunctional PGP subset, highlighting promising candidates for integrated biocontrol strategies. These findings reveal the kiwifruit phyllosphere as a reservoir of functionally diverse and copper-resilient bacteria, highlighting both ecological risks and opportunities for microbiome-based biocontrol innovation. This work establishes a functional baseline for future *in planta* validation and microbiome-informed disease management strategies.

## Introduction

The cultivation of kiwifruit (*Actinidia* spp.) in Portugal dates back to the 1970s, when the ‘Hayward’ variety (*A. chinensis* var. deliciosa) adapted well to local conditions [1]. Encouraged by a favourable climate and a growing market, production grew steadily in the 1980s and 1990s, particularly in the northern and central regions. By the early 2000s, national yields had surpassed 34,000 tonnes per year, placing Portugal among Europe’s key producers [2]. According to C projections, output may increase by another 10 %, reaching around 58,000 tonnes [3]. Today, the crop is mainly grown in Entre Douro e Minho, Beira Litoral, and parts of Beira Interior and the Tagus Valley, where edaphoclimatic conditions are favourable for kiwifruit growth [4].

Kiwifruit was seen as a relatively problem-free crop, with fungal pathogens such as *Botrytis cinerea* and *Diaporthe actinidiae* being the main concerns [5]. However, this changed dramatically following the spread of bacterial canker, caused by *Pseudomonas syringae* pv. *actinidiae* (Psa). Psa causes characteristic symptoms, including leaf spots, brown-black cankers with reddish exudates, flower necrosis, and dieback of shoots and leaders, often leading to severe yield losses and reduced vine longevity. Psa was first reported in Europe in 1992, but the severe outbreaks that turned it into a global threat began in Italy in 2008 [6]. It was detected in Portugal two years later, initially in orchards located in Santa Maria da Feira and Valença [7,8]. The pathogen quickly spread across northern and central regions, severely impacting orchards in the regions of Entre Douro e Minho and Beira Litoral.

Figueira et al. (2020) [9] analysed 604 Psa isolates from four orchards under different abiotic conditions and found substantial genetic diversification since the initial clonal outbreaks in 2010. This diversity varied according to the orchard, season and leaf niche, with a sharp decline in the fall, when a few dominant genotypes prevailed - probably adapted to environmental and management conditions. This persistence highlights the adaptability of Psa and reinforces the need for integrated management strategies adapted to the site. In this context, microbiome-based approaches are emerging as new avenues for sustainable and resilient disease control.

Since the earliest outbreaks of Psa, kiwifruit growers have relied on integrated strategies to manage the disease, combining orchard hygiene, pruning care, and irrigation control with chemical treatments and, where available, tolerant cultivars [6,10]. Copper-based bactericides are the basis in most regions, largely due to their broad activity and relatively low cost. In some countries, antibiotics, such as streptomycin, are also used; however, this is not the case in the EU, where such products are banned due to concerns about antimicrobial resistance [11].

The problem is that copper treatments are not always effective, particularly under high disease pressure. Repeated spraying may contribute to residues on plant tissues and exert selective pressure on bacterial populations [12,13]. Resistance is no longer just a concern for the future: studies from Greece have already identified Psa strains capable of tolerating copper concentrations up to 400 µg/mL [13]. Moreover, resistance genes like *copABCD* have been detected in isolates from both Italy and New Zealand [12,14]. These copper-tolerant Psa isolates were identified in orchards subjected to repeated copper sprays over multiple seasons, highlighting the selective pressure imposed by long-term copper-based management. Within the EU, growers are mostly left with copper and plant defence inducers like acibenzolar-S-methyl [15–19]. These tools can reduce symptoms but rarely control the pathogen. The pressure for alternatives is mounting. Policies like the European Green Deal and the Farm-to-Fork Strategy call for a 50% pesticide reduction by 2030 and promote biological solutions as a way to protect food systems, ecosystems, and biodiversity [20–21]. In that context, understanding copper tolerance locally—including in Portuguese orchards—is not just useful, it’s necessary. Importantly, no copper-resistant Psa strains had been documented in Portugal prior to this study. Given the long-term use of copper-based sprays in Portuguese orchards, assessing whether copper tolerance may already have emerged locally—either in Psa or in co-occurring phyllosphere bacteria that can act as reservoirs of tolerance genes—was therefore particularly relevant.

In recent years, there has been a real shift in how we understand the plant microbiome, now recognised as a living, responsive ecosystem—one that plays active roles in nutrient cycling, stress buffering, disease suppression and immune regulation [22–24]. The focus has moved away from simply replacing agrochemical inputs with biological ones. Instead, the aim is to manage this complex microbial community as a functional part of the plant itself.

New technologies have helped open that door. Synthetic microbial communities (SynComs), microbiome engineering, and microbiome-assisted plant breeding are all being explored as ways to design more stable and site-specific solutions. In practical terms, microbial consortia adapted to local environmental conditions may offer better results than single-strain applications, particularly in open-field systems [23,24].

Kiwifruit is a good example of where this shift matters. Recent works have shown that the phyllosphere microbial communities can play a protective role, helping to buffer disease pressure from pathogens like Psa [25,26]. Studies by Correia et al. [27] and Patterson et al. [28] looked at how these communities respond to different production systems, though their findings are mostly based on culture-independent methods. As a result, we still know little about the functional capabilities of the strains we can actually grow, test, and potentially apply. That gap becomes even more relevant if we consider the plant as a holobiont—a partnership between the host and its microbiota [29,30]. In this frame, pathogens can destabilise the microbial balance, triggering shifts in community composition that may either support recovery or haste plant decline [31,32]. Understanding these shifts is key to boosting microbial allies that can persist under pressure and promote resilience from within.

To address this gap, the present study aimed to explore the culturable bacterial microbiota associated with the leaves of *A. chinensis* var. *deliciosa* in organic orchards affected by Psa infection in Northwest Portugal. Specifically, we aimed to: (i) isolate and identify epiphytic and endophytic bacterial strains from healthy and diseased male and female plants; (ii) provide a preliminary description of the structural diversity of the culturable bacteriome concerning plant health status and sex; (iii) conduct an initial screening of representative isolates for plant growth-promoting (PGP) traits; (iv) assess their antagonistic activity against Psa strains; and (v) evaluate copper tolerance among bacterial isolates. By integrating taxonomic, ecological and functional insights, we further explore how environmental filtering and microbial plasticity shape the resilience of plant-associated communities under biotic stress. Altogether, this constitutes an *in vitro* functional screening framework that lays the foundation for selecting candidate strains for subsequent in planta and field validation. As such, the present work should be viewed as a preliminary functional triage, designed to identify robust native taxa that warrant further testing under greenhouse and orchard conditions.

Through the isolation, functional characterisation, and preservation of native culturable bacteria, this work contributes to expanding the microbial resource base available for future research on microbiome-based approaches for plant health. By targeting bacteria naturally adapted to the kiwifruit phyllosphere, this study provides a starting point for the development of site-specific microbial resources with potential relevance for sustainable disease management strategies.

## MATERIALS AND METHODS

### Sampling and Processing

Sampling was carried out to obtain comparable phyllosphere bacterial communities from healthy and diseased male and female plants under field conditions. Composite sampling was applied to minimise within-plant variability and to ensure that each sample represented a consistent community profile.

This study was conducted in two neighbouring organic kiwifruit orchards located in Northwest Portugal (near Vila Verde - N 41° 68.930008’; W 8° 4079839’), one classified as healthy and the other as diseased [33] based on prior screening for *Pseudomonas syringae* pv. *actinidiae* (Psa) following European and Mediterranean Plant Protection Organisation (EPPO) standards [34]. Diseased plants showed characteristic bacterial canker symptoms, and Psa infection was confirmed by molecular detection according to the EPPO protocol.

Leaves from five *Actinidia chinensis* cv. *deliciosa* ‘Hayward’ and five cv. ‘Tomuri’ plants were collected from the healthy orchard (H, Psa not detected) and the diseased orchard (D, Psa detected). For each plant, ten leaves were sampled from various canopy positions and pooled to form four composite samples: Healthy Males (HM), Healthy Females (HF), Diseased Males (DM), and Diseased Females (DF). Because sex and cultivar correspond in this production system — females being ‘Hayward’ and males being ‘Tomuri’ — the four composite categories (HF, HM, DF, DM) inherently reflect both sex and cultivar.

All samples were collected on the same day (post-bloom, June) to minimise temporal variability in the phyllosphere microbiota. Samples were placed in sterile plastic bags, stored at 4 °C, and processed on the same day. Approximately 150 g of leaves from each composite sample were cut into small pieces and blended with 600 mL of sterile ultrapure water. The resulting extract was filtered through sterile gauze to remove debris and used for isolating epiphytic and endophytic microorganisms through culture-dependent methods, and for independent culture methods as previously described [33].

### Bacterial Isolation and DNA Extraction

The isolation workflow was designed to maximise recovery of the culturable heterotrophic fraction of the phyllosphere across a broad range of physiological niches, thereby enabling downstream functional and taxonomic comparisons between plant conditions. To maximise recovery of facultative aerobic heterotrophic bacteria, serial dilutions (10⁰ to 10⁻⁶) of the plant extract were prepared. From each dilution, 100 µL was plated onto Alkaliphilic Buffered Medium 2 (ABM2) [35] and Reasoner’s 2A agar (R2A) [36], each buffered to pH 5.5, 7.0 and 8.5 at a final concentration of 75 mM. Plates were incubated at 20 °C, 25 °C, and 37 °C. ABM2 and R2A were selected based on their contrasting nutrient content (high and low, respectively). Technical replication was performed for each dilution and condition to ensure robustness of representative culturable diversity. After five days of incubation, the Colony-Forming Units (CFUs) were counted. Colonies were characterised based on morphology (pigmentation, shape, margin, size, and texture) and sub-cultured until pure cultures were obtained. Isolates were cryopreserved in the isolation medium supplemented with 15 % (w/v) glycerol and stored at - 80 °C until further use. Each isolate was assigned an ID code (KWT001) for identification purposes during this study. Total DNA was extracted via thermal shock as previously described [9]. Briefly, a bacterial suspension was prepared in lysis buffer (2 % Tween 20 in 0.1 M NaOH), followed by heat denaturation and sedimentation of cell debris. DNA concentrations were normalised using a Nanodrop spectrophotometer.

### Fingerprinting Analysis

Random Amplified Polymorphic DNA (RAPD) profiling was used to reduce redundancy among isolates and to identify unique genomic fingerprints, ensuring efficient selection of representative strains for sequencing and functional screening.

Bacterial isolates were fingerprinted by RAPD analysis using the primer OPA-03 (5′-AGTCAGCCAC-3′) as described by Costa et al. (2005) [37]. PCR products were separated by electrophoresis on 2% (w/v) agarose gels in 1× TAE buffer, stained with GreenSafe Premium (NZYTech, Portugal), and visualised with a DocTM XR+ imaging system (Bio-Rad, Portugal). Gel images were processed and normalised using BioNumerics software, following procedures detailed in Figueira et al. (2020) [9]. Isolates were clustered based on band number, similarity, and migration patterns relative to a molecular weight marker (NZYDNA Ladder III, NZYTech, Portugal).

### Phylogenetic Analysis

Phylogenetic identification of representative isolates allowed taxonomic resolution of RAPD clusters and enabled assessment of shifts in major bacterial lineages associated with plant health status and sex.

Isolates were first grouped into RAPD profiles. Representative strains from each RAPD cluster were selected for 16S rRNA gene sequencing; for clusters with three isolates, one strain was selected; for clusters with four or more isolates, two strains were selected. 16S rRNA gene fragments (∼1500 bp) were amplified using primers 27F (5′-AGAGTTTGATCCTGGCTCAG-3′) and 1525R (5′-GGTTACCTTGTTACGACTT-3′) [38]. PCR products were purified and sequenced by StabVida (Portugal), using primer 519R (5′-GWATTACCGCGGCKGCTG-3′) [39]. Sequences were quality-checked and edited in UGENE [40]. Sequences were clustered into species-level groups at ≥98 % similarity using CD-HIT Suite [41], and taxonomic identities were assigned through BLAST comparison against the NCBI GenBank database. In cases where the species-level assignment was ambiguous or required confirmation, phylogenetic trees were constructed in ARB software [42] using the neighbor-joining algorithm (NJ) with Jukes-Cantor correction and 1000-bootstrap resampling.

To confirm Psa identification among *Pseudomonas* spp. isolates, specific testing was performed by the Laboratory for Phytopathology from Instituto Pedro Nunes (Portugal) according to EPPO standards [34].

### Bacterial Plant Growth-Promoting Potential

These assays were selected to characterise core functional traits relevant to plant performance and to identify multifunctional bacterial isolates with potential agronomic relevance in kiwifruit systems.

The plant growth-promoting potential of 135 isolates was assessed according to Martins et al. [38], including evaluation of phosphate solubilisation, siderophore production, ammonia production, and Indole-3-Acetic Acid (IAA) production. All tests were performed in triplicate. Qualitative phosphate solubilisation was evaluated following Almoneafy et al. [43]. Isolates were grown on GYA medium [GY broth plus 15 g/L agar], and phosphate solubilisation was indicated by clear halos around colonies after seven days at 28 °C. Siderophore production was assessed using chrome azurol S (CAS) agar plates, with the detection of orange halos after three days at 28 °C, following Almoneafy et al. [43]. Ammonia production was evaluated as described by Singh and Kumar [44], based on colour change (yellow to brown) upon the addition of Nessler’s reagent to peptone water cultures after four days at 30 °C. Results were categorised into three levels: none (0), low (1), and high (2) production. IAA production was quantified using Salkowski’s reagent and colorimetric measurement at 530 nm, following Almoneafy et al. [43]. Isolates were cultured in LB medium supplemented with L-tryptophan (40 µg/mL) at 30 °C, 160 rpm for 48 h. IAA production was analysed by one-way ANOVA followed by Tukey’s multiple comparisons test (p < 0.05), using GraphPad Prism v8.4.3 (San Diego, California, USA).

### Antagonistic Activity against Pseudomonas syringae pv. actinidiae

Antagonism assays were performed to determine whether native phyllosphere bacteria could inhibit Psa growth, providing insight into their potential role in natural disease suppression and biocontrol.

The antagonistic activity of 54 isolates against Psa strain CFBP7286 (Biovar 3) was evaluated using three methods: cross-streak, disk diffusion, and agar well diffusion. Overnight cultures of isolates and Psa were prepared in LB broth at 28 °C. All tests were performed in triplicate, and all three assays were applied to the same set of isolates. Antagonistic activity was analysed by one-way ANOVA followed by Tukey’s multiple comparisons test (p < 0.05), using GraphPad Prism v8.4.3 (San Diego, California, USA).

In the Cross-Streak Method, following Lertcanawanichakul and Sawangnop [45], Psa was streaked at the centre of LB agar plates, and isolates were streaked perpendicularly. Then the inoculated plates were incubated for six days at 28 °C.

In the Single Disk Method, following Lertcanawanichakul and Sawangnop [45], LB agar plates were seeded with Psa suspension and filter discs impregnated with isolates were placed onto the surface. Inhibition zones were measured after six days at 28 °C.

In the Agar Well Diffusion Method, following Tontou et al. (2016) [46], bacterial inocula (10^8^ CFU/mL) were spotted onto LB agar plates and incubated for 48 h. Plates were then sprayed with Psa suspension (10^6^ CFU/mL) and re-incubated for 48 h. The average inhibition area (AIA) was calculated: AIA = π(R² − r²), where R is the radius of the inhibition zone and r is the radius of the colony.

### Copper Tolerance

Copper tolerance was assessed to evaluate the capacity of phyllosphere taxa to withstand copper-based phytosanitary treatments, a key selective pressure in kiwifruit orchards and relevant for interpreting ecological resilience and biocontrol potential.

Copper tolerance was assessed on mannitol–glutamate yeast extract (MGY) agar following the plating procedure of Tontou et al. (2016) [46], with adaptations. Isolates were spotted on MGY supplemented with copper (II) sulphate pentahydrate (CuSO₄·5H₂O) at concentrations of 0.0, 0.8, 1.2, 2.0 and 2.8 mM and incubated at 25 °C for 96 h. The minimum inhibitory concentration (MIC) was defined as the lowest copper concentration at which no visible growth was observed. The 0.8 mM threshold used to classify isolates as copper-resistant follows the MIC boundary commonly adopted in studies of phyllosphere and Psa-associated bacteria, where it represents the established cut-off distinguishing baseline sensitivity from adaptive copper tolerance. In line with this criterion, isolates able to grow at ≥0.8 mM were considered copper-resistant [13,47]. Plates without copper served as growth controls, and all assays were performed in duplicate.

## RESULTS

### Bacterial Community Structure Based on Culturable Isolates

To characterise the culturable phyllosphere bacteriome of *Actinidia chinensis* under Psa pressure, the taxonomic composition of bacterial isolates obtained from healthy and diseased male and female plants was examined. The culturable bacterial community isolated from kiwifruit leaves exhibited differences in structure between healthy and diseased plants and between male and female plants. In total, 1058 strains were isolated and grouped into 261 RAPD clusters and assigned to 58 species from 29 genera. RAPD clustering showed high correspondence with 16S-based identification, supporting the validity of this grouping strategy. Figure 1 summarises the relative abundance of the recovered genera across the different sample types, highlighting both shared and condition-specific taxa. Five genera — *Curtobacterium*, *Microbacterium*, *Pantoea*, *Pseudomonas*, and *Sphingomonas* — were consistently detected across all sample types, representing the core culturable bacteriome of *A. chinensis* leaves. Among these, *Pseudomonas* and *Pantoea* were the most abundant genera across diseased plants (DF and DM), where *Pseudomonas* reached up to 56.5 % and *Pantoea* up to 36.1 %. In contrast, healthy plants exhibited a more even distribution, with *Actinomycetota* genera such as *Curtobacterium*, *Microbacterium* and *Frigoribacterium* contributing substantially to community composition. Beyond the dominant taxa, additional shifts in the relative representation of several genera were observed across plant health status and sex. *Sphingomonas* and *Erwinia* tended to show higher relative abundances in diseased plants, whereas Enterobacter and *Pluralibacter* were proportionally more represented in female plants under both health conditions. In addition, *Frigoribacterium* was particularly prominent in healthy female plants, contributing markedly to their overall community composition (Figure 1).

**Figure 1.**
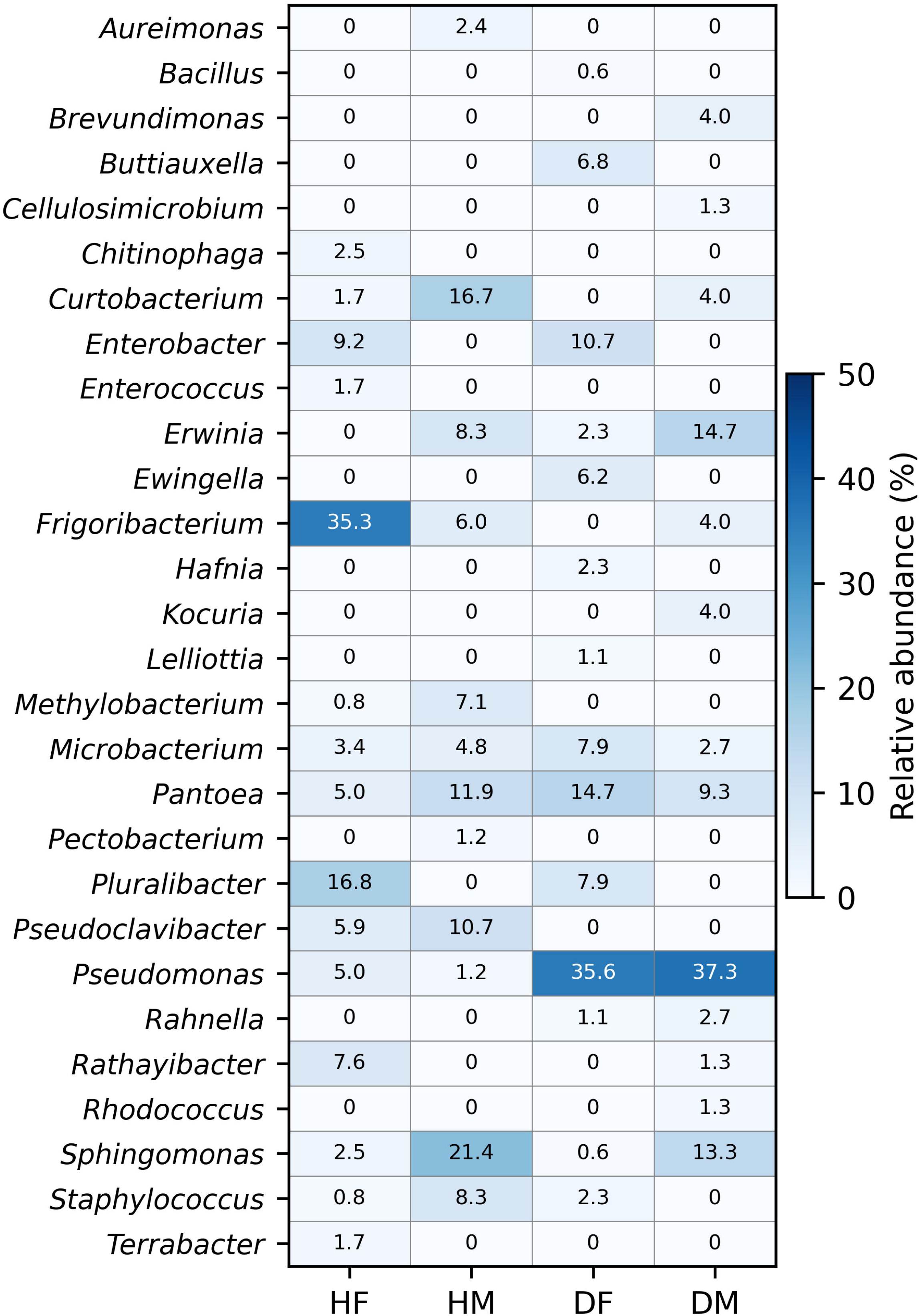
Relative abundance (%) of bacterial genera isolated from the phyllosphere of *Actinidia chinensis* var. *deliciosa* under four plant conditions: Healthy Female (HF), Healthy Male (HM), Diseased Female (DF), and Diseased Male (DM). Data are based on culture-dependent identification of 1,058 isolates grouped into genera. Colour intensity indicates the proportion of each genus in the respective sample types. Only genera with relative abundance ≥0.5 % in at least one sample type are shown (Supplementary Table S1). The data highlights the dominance of *Pseudomonas* in diseased plants and the higher representation of *Actinomycota*-associated genera such as *Curtobacterium* and *Frigoribacterium* in healthy plants.

### Phylogenetic Insights into the Culturable Bacteriome

To assess how major bacterial lineages were distributed across plant conditions, the phylogenetic composition of the representative isolates assigned to RAPD clusters was examined. Two hundred and eighteen representative isolates from RAPD clusters were sequenced for the 16S rRNA gene [∼1,500 bp], resulting in the identification of 58 species distributed across 29 genera (Supplementary Table S1). Most species belonged to the phylum *Pseudomonadota*, particularly in diseased plants, where they accounted for 96.1 % of isolates in DF and 87.0 % in DM (Figure 1). The genus *Pseudomonas* was particularly abundant in both diseased groups, reflecting its dominance under canker pressure.

In contrast, healthy leaves exhibited a more phylogenetically diverse community, with *Actinomycetota* representing 28.2 % and 45.1 % of the isolates in HF and HM, respectively. Genera such as *Curtobacterium*, *Microbacterium* and *Frigoribacterium* were notably more frequent in healthy plants than in diseased ones. Minor proportions of *Bacillota* and *Bacteroidota* were also detected, the latter being exclusive to healthy female plants, suggesting that some low-abundance groups may be sensitive to disease-associated shifts in community structure.

### Distribution of Multifunctional Plant Growth Promoting Bacteria and Copper Tolerance

From the 135 bacterial isolates screened for plant growth-promoting (PGP) traits—indole-3-acetic acid (IAA) production, siderophore production, ammonia production, and phosphate solubilisation— and copper tolerance, 25 isolates exhibited all four PGP traits and were classified as multifunctional candidates. These isolates were recovered from all four plant conditions, with a higher proportion originating from DM and HF plants (Figure 2A, Table S2), indicating that multifunctionality is not restricted to healthy tissues and can persist under disease pressure.

**Figure 2.**
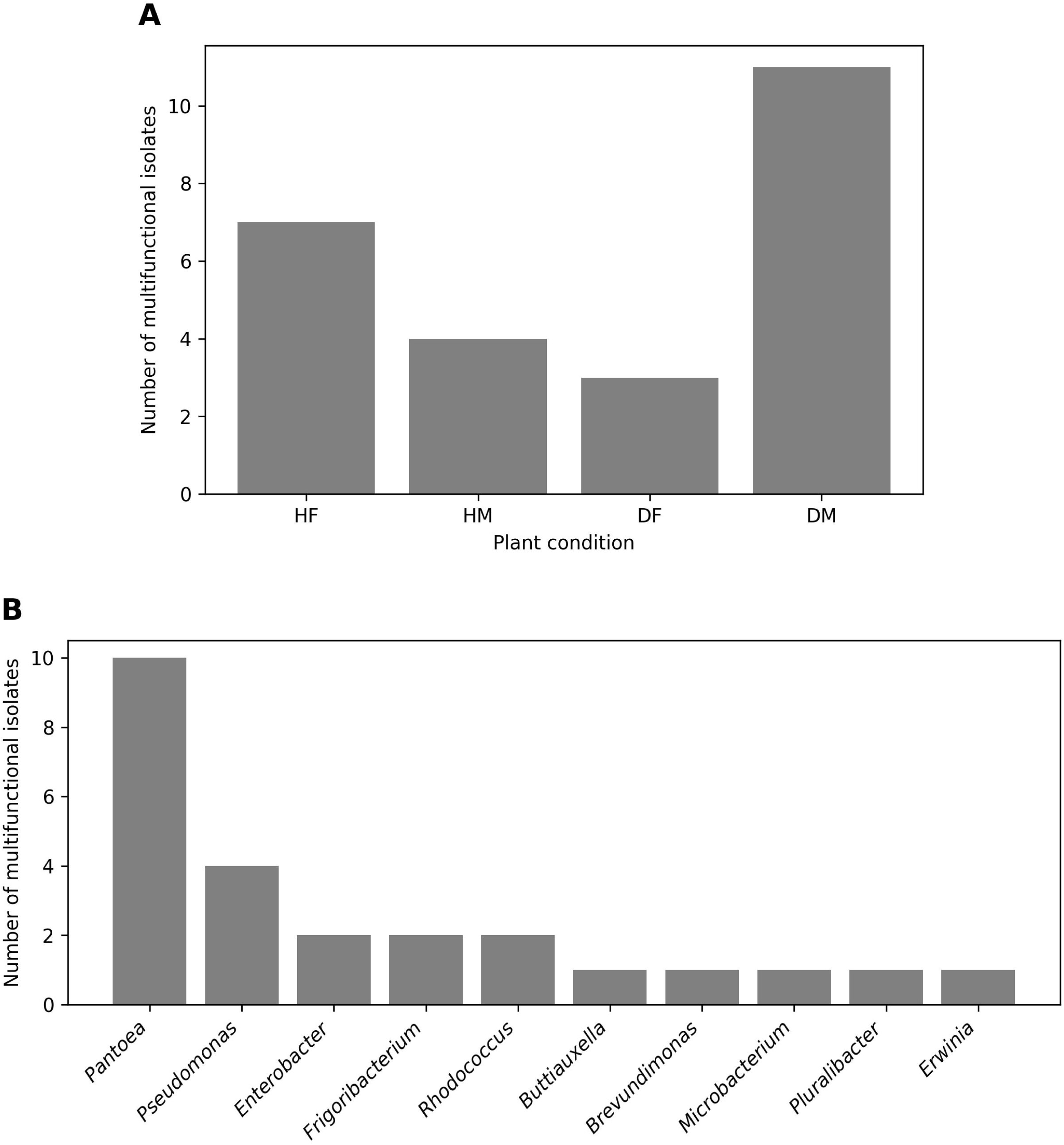
Distribution and taxonomic composition of multifunctional plant growth-promoting bacteria isolated from the kiwifruit phyllosphere. **(A)** Number of bacterial isolates exhibiting all four evaluated plant growth-promoting (PGP) traits (indole-3-acetic acid production, siderophore production, phosphate solubilisation and ammonia production) recovered from each plant condition: healthy female (HF), healthy male (HM), diseased female (DF) and diseased male (DM) plants. **(B)** Number of multifunctional isolates per bacterial genus, showing a strong concentration of multifunctionality in a limited number of taxa. Genera represented by a single isolate are shown individually. Detailed strain-level functional profiles are provided in Table S2.

Analysis of the taxonomic distribution of the multifunctional isolates revealed a strong concentration in a limited number of genera (Figure 2B). *Pantoea* and *Pseudomonas* together accounted for more than half of the multifunctional strains. Additional multifunctional isolates belonged to *Enterobacter*, *Frigoribacterium* and *Rhodococcus*, whereas the remaining genera (*Buttiauxella*, *Brevundimonas*, *Microbacterium*, *Pluralibacter* and *Erwinia*) were represented by a single isolate each, indicating a highly uneven distribution of multifunctionality across taxa. Notably, several genera that were abundant in the broader culturable community (e.g. *Sphingomonas* and *Curtobacterium*) were not represented among the multifunctional isolates, highlighting that numerical dominance does not necessarily translate into functional dominance.

Copper tolerance was widespread among the culturable phyllosphere community. Using the 0.8 mM cut-off commonly applied to phyllosphere bacteria associated with Psa [13,47], 46.7 % (63/135) were classified as copper-sensitive (MIC < 0.8 mM), while 53.3 % (72/135) were resistant (MIC ≥ 0.8 mM) (Table S2). Most resistant isolates displayed MIC values of 0.8–1.2 mM, although a small number reached 2.0 mM, indicating substantial tolerance within the phyllosphere community.

Several non-pathogenic taxa exhibited combinations of plant growth-promoting traits and copper tolerance, as revealed by the functional screening (Table S2). These included representatives of *Pantoea agglomerans* (KWT1371, KWT1372, KWT1355), *Enterobacter asburiae* (KWT1367) and *Erwinia iniecta* (KWT9), which maintained relevant functional traits under copper exposure. Importantly, multifunctionality and copper tolerance were not strictly associated, as several multifunctional isolates were copper-sensitive, highlighting the independence of these functional dimensions within the phyllosphere community.

### Antagonistic Activity against Pseudomonas syringae pv. actinidiae and the Potential Application of Bacillus pumilus in Kiwifruit Disease Management

To determine whether native phyllosphere bacteria could suppress Psa, the antagonistic potential of 54 isolates was assessed using three complementary inhibition assays. Overall, approximately 10 % of the isolates inhibited Psa in at least one method, indicating that antagonistic capacity is widespread within the culturable phyllosphere microbiota. In the cross-streak assay, two isolates, *Pseudoclavibacter helvolus* (KWT786) and *Bacillus pumilus* (KWT1381), produced visible growth inhibition against Psa, typically manifesting as narrow inhibition zones along the streak interface. The disk diffusion assay confirmed antimicrobial activity in a subset of isolates, which produced inhibition halos with an average diameter of 0.94 ± 0.12 mm. The agar well diffusion method provided quantitative estimates of antagonistic strength, with Average Inhibition Area (AIA) values ranging from 221.19 to 228.17 mm², and several isolates displaying strong inhibition comparable to known biocontrol agents.

Among the tested strains, *B. pumilus* (KWT1381) consistently displayed strong antagonism across all three methods. In the agar well diffusion assay, this isolate produced a clear inhibition halo measuring 22 mm in diameter, corresponding to an AIA of 228.17 mm². Notably, some potential antagonist isolates overlapped with the multifunctional subset described in Section 2.3, indicating that plant growth-promoting traits and pathogen suppression may co-occur in certain native strains, thereby highlighting particularly promising native candidates for integrated biocontrol strategies.

## DISCUSSION

Relative abundance analysis showed that the *Pseudomonas* genus dominated the culturable bacteriome in diseased plants, reaching more than half of the total isolates in diseased plants. In contrast, healthy samples displayed a more balanced distribution among genera, with *Frigoribacterium* and *Curtobacterium* contributing significantly to the community structure. Overall, disease presence was associated with a marked decrease in community evenness and diversity, favouring the dominance of opportunistic genera such as *Pseudomonas* and *Pantoea*. These results are consistent with the Anna Karenina Principle in microbial ecology, which posits that stressed hosts tend to harbour more variable and less even microbial communities [32,48]. Dysbiosis of the phyllosphere microbiota under biotic stress has been described in several plant systems [31], including kiwifruit [25–27]. Our findings complement these studies by providing a culture-based perspective on this broader pattern.

The timing of sampling (late spring) was particularly relevant, as in Portugal, Psa populations expand during spring under favourable temperatures (12–18 °C) and high humidity, while consecutive summer heat days lead to a sharp reduction in diversity and dominance of a few adapted clones [9]. Capturing the microbiota during this peak of pathogen activity provided a representative snapshot of microbiome shifts under strong infection pressure.

To contextualise our findings, the isolated bacteriome in this study was compared with the phyllosphere microbiome previously characterised by metabarcoding using the same plant material [33]. Despite methodological differences, both datasets identified *Pseudomonadota* as the dominant phylum, with *Pseudomonas* and *Sphingomonas* as key genera. Our culture-based approach recovered several taxa also reported by Ares et al. (2021) [33], including *Curtobacterium*, *Frigoribacterium*, *Microbacterium*, and *Methylobacterium*. In contrast, metabarcoding captured a broader diversity, including *Hymenobacter* and *Massilia*, reflecting the intrinsic bias of culture-based recovery towards fast-growing or nutritionally versatile bacteria.

Our results complement these studies by providing a culture-based inventory of experimentally validated strains with plant-beneficial traits, thus bridging the gap between sequence-based inference and applied microbial selection. Unlike previous work that relied mainly on predictive functional profiling, such as Correia et al. (2025) [27] and Patterson et al. (2025) [28], the present collection offers concrete candidates for testing as potential biocontrol agents under biotic stress conditions. Importantly, this work goes beyond descriptive cataloguing by incorporating a systematic functional screening, allowing the identification of concrete native strains with validated agronomic potential.

This convergence with other culture-independent surveys of Actinidia phyllosphere [25–27, 49] reinforces the view that Psa infection strongly reduces diversity and favours opportunistic taxa such as *Pseudomonas*. Genera such as *Sphingomonas* and *Methylobacterium*, consistently detected and proportionally more represented in healthy plants across both approaches, stand out as putative contributors to resilience. Certain *Sphingomonas* strains, such as *S. sediminicola* Dae20, have been shown to enhance systemic resistance against foliar pathogens through the activation of jasmonic acid and ethylene signalling, highlighting their role in promoting plant immunity and potentially stabilising phyllosphere microbiota under stress [50,51]. Similarly, *Methylobacterium* spp. are known to synthesise phytohormones such as cytokinins, contribute to plant stress alleviation, and exhibit biocontrol activity against several pathogens [52].

Overall, these findings suggest that Psa infection acts as a strong environmental filter, restructuring the phyllosphere microbiome in line with the holobiont framework [30,53]. Integrating culture-dependent and molecular approaches, therefore, provides complementary perspectives and helps identify robust microbial candidates for sustainable disease control in kiwifruit. This culture-based validation also resonates with recent perspectives emphasising that preserving and harnessing microbial diversity is central to future plant health strategies, positioning native strain collections as key resources for biocontrol innovation [22].

Regarding the cultivable bacteriome described in item 2.2, the results are consistent with previous studies showing that Psa infection reshapes phyllosphere microbial structure by reducing diversity and favouring *Pseudomonadota* dominance [25–27,33].

At the genus level, *Pseudomonas*, *Curtobacterium*, *Microbacterium*, *Pantoea*, and *Sphingomonas* were recovered from all sample types, forming the core culturable microbiota of *A. chinensis*. These genera have also been consistently reported in culture-independent surveys [33], reinforcing their ecological relevance. The persistent presence of *Sphingomonas* and *Methylobacterium* in healthy plants suggests their potential roles in plant protection and microbiome stability, given their known capacity to modulate plant immunity, produce antimicrobial compounds, and enhance plant growth under stress [50–52,54].

Several genera were restricted to a single sample type. For example, *Terrabacter* and *Enterococcus* were exclusive to HF, *Geodermatophilus* and *Dermacoccus* to HM, and *Raoultella*, *Hafnia*, and *Pectobacterium* to DF. This pattern indicates that both disease status and plant sex shape the culturable microbiota, although pathogen pressure appears to be the main driver [33,53]. Such selective pressure may destabilise the microbiome assemblies, leading to dysbiosis, consistent with models proposing that loss of microbial homeostasis compromises plant resilience [31]. Within the holobiont framework, this disruption extends beyond the plant host, affecting the entire plant–microbiota system [30]. Moreover, although some sex-related differences (HF vs. HM) were detected, disease status was the dominant factor shaping bacterial communities, consistent with host-mediated filtering processes [53].

Although sampling depth was not sufficient to capture the full taxonomic diversity, particularly in diseased plants, the combined RAPD and 16S profiling was adequate to reveal the dominant trends and taxonomic shifts across conditions. We acknowledge that relying solely on 16S rRNA sequencing limited species-level resolution for a subset of isolates; future work will apply higher-resolution markers to refine taxonomic assignments and strengthen ecological interpretations. These results align well with patterns described in culture-independent surveys, providing complementary culture-based insights into the effects of Psa infection on phyllosphere communities.

The analysis of the distribution of multifunctional PGPBs revealed that these isolates were not restricted to healthy environments and could persist or even proliferate under disease pressure, particularly in male plants. At the genus level, multifunctionality was strongly concentrated in *Pantoea* and *Pseudomonas*, which together accounted for more than half of the multifunctional isolates. Both genera were also widely distributed across all sample types in the broader culturable dataset, reinforcing their ecological relevance within the kiwifruit phyllosphere. Their repeated representation among multifunctional isolates suggests that ecological success in this system is associated with functional versatility, particularly under contrasting host health conditions. Additional multifunctional isolates belonged to *Enterobacter*, *Frigoribacterium* and *Rhodococcus*, indicating that multifunctionality, although unevenly distributed, spans phylogenetically diverse taxa.

Interestingly, *Enterobacter* (including multifunctional strains) and *Klebsiella* (within the broader PGPB set) - both affiliated with *Enterobacteriaceae* family - were identified among PGPB isolates from both healthy and diseased plants. Their functional versatility, coupled with their presence in copper-tolerant groups (described in section 2.3), underscores their adaptability to contrasting environmental conditions. By contrast, *Sphingomonas* and *Methylobacterium* dominated healthy samples in terms of abundance but were not represented among the multifunctional isolates, whereas *Frigoribacterium* was represented by a limited number of multifunctional strains. This suggests that numerical prevalence does not necessarily correspond to functional dominance. Such ecological-functional mapping highlights the resilience and potential of specific genera as bioinoculants in Actinidia cultivation, while also suggesting that diseased plants, particularly males, may act as reservoirs of robust taxa selected under biotic and abiotic stress.

Notably, one Psa isolate tolerated copper up to 2.0 mM, matching the upper resistance levels reported for Greek Psa field populations [13]. To our knowledge, this constitutes the first evidence of highly copper-resistant Psa in Portuguese orchards.

Remarkably, *Erwinia iniecta* (KWT9) displayed the highest resistance detected, with growth up to 2.8 mM (Table S2). This surpasses thresholds previously reported for Psa in Greek orchards (2.0 mM) [13] and exceeds values described for other phyllosphere-associated bacteria [47]. Such exceptional resistance in a non-pathogenic taxon highlights the potential of the kiwifruit phyllosphere to act as a reservoir of extreme copper resistance.

The recurrence of *Pantoea agglomerans* and *Curtobacterium flaccumfaciens* across both healthy and diseased plants highlights their ecological versatility, while *Rahnella inusitata* and *Enterobacter asburiae* were consistently associated with healthy plants, supporting their potential role as native bioinoculants. Their presence in both healthy and diseased plants—particularly in male plants—suggests that copper-managed orchards may act as reservoirs of functionally robust taxa capable of persisting under chemical selection pressure.

In comparative terms, the proportion of copper-resistant isolates recovered in this study (53.3 %) is strikingly higher than values reported in previous work on plant-associated bacteria. Beltrán et al. (2021) [47] found that only 10–20 % of *Pseudomonas* isolates from stone fruit phyllosphere in Chile were resistant at the 0.8 mM threshold, with the majority classified as sensitive. Similarly, Thomidis et al. (2025) [13] reported copper resistance in Greek Psa populations as a subset phenomenon, with resistant isolates reaching up to 2.0 mM but never representing the majority. Other studies in phytopathogenic bacteria, such as *Xanthomonas euvesicatoria* [55], likewise described resistance as a minority trait at the field level. In contrast, our results indicate that copper resistance is already predominant in the Portuguese kiwifruit phyllosphere, encompassing both pathogenic and non-pathogenic taxa.

Recent evidence confirms the increasing prevalence of copper-resistant Psa strains in European orchards, particularly in Italy and Greece, with minimum inhibitory concentrations [MICs] reaching or exceeding 1.2 mM, and in some cases up to 2.0 mM [12; 13]. Although no published studies have yet reported copper-resistant Psa strains from Portugal, the selective pressure imposed by copper-based treatments in local orchards may favour their emergence. In this context, the identification of native non-pathogenic and multifunctional strains, including *E. iniecta*, which grew at concentrations exceeding 2.0 mM, reaching up to 2.8 mM in the case of strain KWT9, while *Curtobacterium flaccumfaciens* tolerated copper at levels up to 2.0 mM (Table S2) - highlights their added value as candidates for integrated management in high-copper environments.

These findings underscore that the kiwifruit phyllosphere acts as a reservoir of beneficial and copper-resilient bacteria adapted to agrochemical stress. The multifunctionality and resilience of these strains may reflect evolutionary responses to plant-derived selection cues, stress factors, and intermicrobial interactions within the phyllosphere. Such traits reinforce the value of culture-dependent approaches for identifying native microbial resources with agronomic potential and support the integration of microbial functional traits into sustainable disease management frameworks.

Genera such as *Pantoea*, *Pseudomonas*, and *Enterobacter* emerge as particularly promising for biotechnological use, combining plant growth promotion, stress resilience, and potential disease suppression. Their presence across different health statuses and plant sexes suggests broad ecological plasticity, making them suitable candidates for microbiome-based interventions in diverse orchard conditions. This supports the idea that microbial adaptability to host environments is a key feature of successful plant-associated taxa [29].

Our findings complement recent studies by Fu et al. (2024) [26] and Correia et al. (2025) [27], which used metagenomic and transcriptomic data to infer microbial responses to Psa infection. While those studies highlighted functional potential, our culture-based methodology provides *in vitro* validation of traits, such as copper tolerance and PGP activities and identifies concrete candidate strains for future in planta testing. This functional confirmation expands the microbial toolbox for tailored microbiome-based interventions. The resulting collection of resilient and functionally diverse strains represents a valuable asset for applied microbiome research.

Finally, the discovery that common phyllosphere bacteria, including well-documented plant associates like *Pantoea agglomerans* but also less-studied taxa such as *Rahnella inusitata* and *Microbacterium testaceum*, act as reservoirs of copper resistance is of significant ecological concern. While most studies on the microbial resistome in agricultural settings have focused on soil [56–58], our results suggest that the selection pressure from copper-based treatments creates a similar genetic pool of resistance in the phyllosphere. Copper resistance in some plant pathogens is well-documented [14], but our findings highlight that the issue extends beyond specific phytopathogenic taxa. The presence of high-level resistance in ubiquitous, non-pathogenic species raises concerns about the potential for horizontal gene transfer, which can facilitate the dissemination of resistance traits to other members of the microbial community, including emerging pathogens [59]. The occurrence of resistance in genera like *Pantoea*, which are well-known for their plant-associated lifestyle and potential for gene exchange, is particularly concerning [47,60]. Consequently, the kiwifruit phyllosphere may serve as a hub for the evolution and spread of copper resistance, with broader implications for sustainable disease management across a range of crops [13]. While these findings highlight the adaptive capacity of phyllosphere communities, the potential application of copper-tolerant strains requires careful consideration. Any future use will necessitate a dedicated biosafety evaluation, including assessments of genetic stability and the potential for horizontal transfer of resistance determinants under field conditions.

The detection of both highly resistant Psa and exceptionally resistant non-pathogenic isolates such as *E. iniecta* underscores the dual ecological role of copper tolerance in the phyllosphere. On the one hand, it facilitates pathogen persistence under chemical management; on the other, it highlights the adaptive capacity of commensal taxa that may be harnessed as biocontrol allies in copper-managed environments. However, copper resistance in Psa has been linked to mobile genetic elements, including integrative conjugative elements (ICEs) and plasmids carrying *copABCD* and *cus/czc*, whose transfer has been demonstrated both in vitro and in planta [14]. This highlights the need for vigilance when deploying copper-tolerant commensals as biocontrol agents [59] and underscores the broader ecological implications of copper resistance in phyllosphere microbial communities.

After evaluating the isolates for antagonistic activity and differences between plant conditions, we observed that *Pseudomonas* and *Pantoea* isolates had a higher proportion of active strains on diseased male plants, while healthy female plants contributed fewer antagonistic isolates. These patterns suggest that pathogen pressure may enrich antagonistic taxa in certain host niches. Notably, potential antagonist isolates also exhibited copper tolerance or overlapped with the multifunctional subset, reinforcing their potential as resilient candidates for integrated disease management strategies.

*B. pumilus* is already employed in commercial biocontrol formulations targeting a range of phytopathogens in crops such as soybean, maize, strawberry, and sugarcane. However, to our knowledge, this is the first report documenting its antagonistic potential against Psa in kiwifruit. This finding highlights the opportunity to broaden the repertoire of biocontrol agents available for Actinidia cultivation and reinforces the importance of culture-based approaches for identifying novel applications of well-characterised microbial strains.

The observed antagonism suggests that *B. pumilus* could be incorporated into integrated disease management strategies for kiwifruit, particularly in scenarios where copper-resistant Psa strains emerge and consumer demand for agrochemical-free solutions intensifies. Although *B. pumilus* KWT1381 did not exhibit copper tolerance, its strong antagonistic effect against Psa supports its potential as a complementary agent in microbiome-based strategies, particularly valuable under reduced-copper or agrochemical-free management systems. These antagonistic potentials complement previous reports of biocontrol against Psa using *Aureobasidium pullulans* or *Actinobacteria* [46,61], situating our findings within a growing portfolio of microbial solutions.

The presence of active strains in both healthy and diseased plants suggests that natural antagonists may contribute to microbiome-mediated suppression of Psa, even under pathogen pressure. Genera such as *Pantoea* and *Pseudomonas* have previously been reported as effective biocontrol agents against diverse phytopathogens, and our results extend their relevance to the kiwifruit–Psa pathosystem. Notably, several antagonistic isolates originated from diseased plants, indicating that pathogen-challenged environments may act as reservoirs of functionally robust strains adapted to stress.

## CONCLUSIONS

This study provides an integrative assessment of the culturable phyllosphere bacteriome of *Actinidia chinensis* under pressure from *Pseudomonas syringae* pv. *actinidiae* (Psa) infection. By combining taxonomic, ecological and functional analyses, we demonstrate that disease status strongly influences microbial structure, reducing diversity and promoting the dominance of opportunistic *Pseudomonadota* such as *Pseudomonas* and *Pantoea*. These compositional shifts align with the Anna Karenina Principle, suggesting that Psa-induced dysbiosis disrupts the holobiont balance of the phyllosphere.

Despite this disruption, a subset of bacterial isolates exhibited multifunctional plant growth-promoting traits, while copper tolerance and in vitro antagonistic activity against Psa were also detected across the culturable collection. Notably, these functional traits can co-occur in native strains, supporting the identification of promising candidates for integrated biocontrol strategies. Among the isolates showing in vitro antagonism, *Bacillus pumilus* (KWT1381) consistently inhibited Psa, broadening the repertoire of genera with biocontrol potential and representing a novel application for this well-known biocontrol agent.

In addition, we report a strikingly high prevalence of copper resistance across the phyllosphere microbiota, with more than half of the isolates classified as resistant at the 0.8 mM cut-off, including both pathogenic and non-pathogenic taxa. We report for the first time in Portugal, a highly copper-resistant Psa isolate (up to 2.0 mM), and an exceptionally resistant non-pathogenic strain of *Erwinia iniecta* (2.8 mM), underscoring both the risk of resistance spread and the opportunity to harness robust native taxa under copper pressure. These findings underscore the strong selective pressure imposed by copper-based treatments and highlight the dual ecological role of copper tolerance: enabling pathogen persistence while also selecting commensal taxa that may be harnessed for biocontrol under copper stress.

Overall, the results highlight the value of culture-dependent approaches in identifying native microbial resources with agronomic potential. As this work is based on *in vitro* functional assays, the findings should be interpreted as an initial screening step rather than direct evidence of field efficacy. Accordingly, future work will prioritise *in planta* validation, the development of synthetic consortia, and field trials under copper-managed conditions to bridge the gap between laboratory screening and effective, sustainable disease management in kiwifruit production. Together, these results provide a functionally validated culture collection that can serve as a foundation for targeted microbiome-based interventions in future kiwifruit disease management.

## ACKNOWLEDGEMENT

We thank the kiwifruit orchard owner of “Delicias do Tojal” for providing the Actinidia samples.

## AUTHOR CONTRIBUTIONS

Conceptualization: JC, IT; Data curation: VC, JC, IT, CC, EG; Formal analysis: VC, JP, CC, EG, AA; Funding acquisition: JC; Investigation: VC, JP, EG, CC, DF, AA; Methodology: AA, VC, EG, CC, DF; Project administration: JC; Resources: JC; Software Supervision: JC, IT; Writing – original draft: VC, JC, IT, CC; Writing – review and editing: all authors. All authors have read and agreed to the published version of the manuscript.

## SUPPLEMENTARY MATERIALS

The following supporting information can be downloaded at: https://www.mdpi.com/article/doi/s1; Table S1: Overview of the culturable phyllosphere bacterial diversity associated with Actinidia chinensis var. deliciosa. The table lists representative isolates from each RAPD cluster, identified to genus and, when possible, species level. For each entry, the RAPD cluster code, isolate ID, and total number of isolates per cluster are indicated. Table S2: Identification and functional traits of the 135 bacterial isolates evaluated in this study. Species were identified based on 16S rRNA sequencing of representative isolates from RAPD clusters. Functional traits assessed include phosphate solubilization, siderophore production, ammonia production (rated on a scale from 0 = none to 2 = high, based on colour intensity), and indole-3-acetic acid (IAA) production, both qualitative (+/–) and quantitative (µg/mL). Copper tolerance was determined as the minimum inhibitory concentration (MIC, mM) of CuSO₄·5H₂O. Isolates with MIC <0.8 mM were classified as copper-sensitive (S), while those with MIC ≥0.8 mM were classified as copper-resistant (R) [13; 47].

